# Omitting end preparation reduces index misassignment in Nanopore-based DNA metabarcoding: application to a decadal coastal time series in the Sea of Okhotsk

**DOI:** 10.64898/2026.09.08.750005

**Authors:** Masatoshi Endo, Satoshi Nagai, Tsuyoshi Watanabe, Noriko Kurita, Shuichi Asakawa, Kazutoshi Yoshitake

## Abstract

Environmental DNA (eDNA) metabarcoding enables sensitive, non-invasive assessment of fish communities, but highly multiplexed analyses using Oxford Nanopore Technologies (ONT) platforms require stringent control of sample-index misassignment and sequencing errors. We developed a MiFish experimental workflow combining unique dual indexes, a library protocol that omitted end preparation and used 5′-phosphorylated primers, BLAST-based demultiplexing, quality-dependent clustering, consensus generation, and haplotype partitioning by SNP/INDEL patterns. Omitting pooled end-prep reduced the mean index-chimera rate from 0.0674% to 0.000420%, a 160-fold reduction. We applied the workflow to an archive of seawater samples collected weekly off Monbetsu, Hokkaido, Japan, from 2012 to 2022. Relative read abundance (RRA) data were obtained for 274 samples, and eight taxa showed significant seasonality. For six of these taxa, the three-year mean RRA peaks coincided with reported spawning periods in Hokkaido. The workflow substantially reduced index misassignment and enabled cost-efficient, highly multiplexed analysis of a decadal fish eDNA time series.

## Introduction

Environmental DNA (eDNA) analysis has transformed aquatic biomonitoring by enabling the detection of organisms from DNA released into water without relying exclusively on capture or visual surveys (Bohmann et al. 2014; Thomsen and Willerslev 2015). DNA metabarcoding extends this approach to community-level assessment by simultaneously amplifying and sequencing a taxonomically informative marker from many organisms (Taberlet et al. 2012). For fishes, the MiFish primer system targeting an approximately 170-bp region of mitochondrial 12S rRNA has become widely used because it combines broad taxonomic coverage with high analytical sensitivity (Miya et al. 2015, 2020).

Robust evaluation of seasonal and interannual changes in fish communities requires high-frequency observations collected consistently over multiple years. The coastal waters of the Sea of Okhotsk off Monbetsu in northern Hokkaido form a subarctic environment characterized by seasonal sea ice, marked changes in water temperature, and the turnover of warm-water, cold-water, coastal, and migratory fishes. An archive of seawater samples collected weekly at a fixed site from 2012 to 2022 therefore provides a rare opportunity to reconstruct long-term fish eDNA dynamics at high temporal resolution. The archive and its original plankton-focused monitoring framework have been described previously (Sildever et al. 2023).

Analysis of a large archive also requires a sequencing system that can process many samples flexibly and at low cost. Oxford Nanopore Technologies (ONT) platforms provide portable, real-time sequencing capabilities and have been used for rapid field-based DNA barcoding (Pomerantz et al. 2018). However, sample-index misassignment is a general concern in multiplexed sequencing. On Illumina platforms, index swapping can occur during library preparation and patterned-flow-cell amplification, whereas tag jumping during metabarcoding can transfer sequences between samples and inflate apparent diversity (Schnell et al. 2015; Costello et al. 2018). Even a small number of misassigned reads can affect the interpretation of rare eDNA detections. In addition, individual ONT reads may contain more base-calling errors than short Illumina reads, making consensus generation especially important for improving sequence accuracy (Baloğlu et al. 2021; Dubois et al. 2024).

This study had two objectives. First, we tested whether replacing pooled end preparation with ligation-compatible ends generated during the second PCR could reduce index misassignment in a highly multiplexed ONT MiFish workflow. We also developed a BLAST-based dual-index demultiplexer and a quality-dependent consensus pipeline that repartitions reads using recurrent SNP/INDEL patterns. Second, we applied the optimized workflow to the Monbetsu archive, evaluated the effect of changes in filtration and DNA-extraction protocols on fish eDNA recovery, and characterized seasonal patterns in the relative read abundance of fish taxa.

## Materials and methods

### Study site and archived samples

Surface seawater was collected at the Okhotsk Tower off Monbetsu, Hokkaido, Japan (44°20′12″N, 143°22′54″E). Sampling was conducted approximately once per week throughout the year, generally around 11:00, from 2012 to 2022 as part of long-term monitoring of marine plankton communities. Archived DNA extracts contained bulk environmental DNA, including fish-derived eDNA (Sildever et al. 2023).

Filtration and extraction protocols changed over the course of the sampling period. From April 2012 to March 2018, water was filtered through Nuclepore membranes with pore sizes of 8 and 1 µm (GE Healthcare); DNA was extracted from each filter using Chelex 100 (Walsh et al. 1991), and the two extracts were combined in equal volumes. From April 2018 to March 2019, only a 1-µm Nuclepore membrane was used, followed by Chelex extraction. From April 2019 onward, water was filtered through a 0.22-µm Sterivex cartridge (Merck KGaA), and DNA was extracted with a DNeasy PowerSoil Kit (QIAGEN). Filtration time was limited to 20 min. Filters were stored at −80°C until extraction, and extracted DNA was stored at −20°C until analysis.

### MiFish 12S amplification and unique dual indexing

Fish community analyses targeted an approximately 170-bp segment of the mitochondrial 12S rRNA gene. MiFish-U, MiFish-Ev2, and MiFish-U2 primers were mixed at a ratio of 2:1:1 for the first PCR (Miya et al. 2015, 2020). The forward and reverse locus-specific primers carried the following 5′ tails for amplification in the second PCR:

~~~
Forward tail: ACACTCTTTCCCTACACGACGCTCTTCCGATCT
Reverse tail: GTGACTGGAGTTCAGACGTGTGCTCTTCCGATCT
~~~

Each 10-µL first-PCR reaction contained 5 µL repliQa HiFi ToughMix (QuantaBio), 1.5 µL of each 2 µM primer mixture (forward and reverse; final concentration, 300 nM each), 1 µL nuclease-free water, and 1 µL template DNA. Nuclease-free water was used in the no-template control. Thermal cycling consisted of 95°C for 3 min; 45 cycles of 98°C for 10 s, 65°C for 5 s, and 68°C for 1 s; and 68°C for 1 min. Amplification was assessed by electrophoresis on a 2% agarose gel.

Unique dual indexes were added to first-PCR products using sample-specific combinations of forward and reverse primers in the second PCR. Sixty forward and sixty reverse index primers were prepared (Table S1). Primer IDs 21–57 contained 10-bp indexes, whereas IDs 1–20 and 58–60 contained 24-bp indexes. The primer architectures were:

~~~
Forward: 5′-AATGATACGGCGACCACCGAGATCTACAC-[index]-
ACACTCTTTCCCTACACGACGCTCTTCCGATCT-3′
Reverse: 5′-CAAGCAGAAGACGGCATACGAGAT-[index]-GTGACTGGAGTTCAGACGTGTGCTCTTCCGATCT-3′
~~~

Each 10-µL second-PCR reaction contained 0.05 µL TaKaRa Ex Taq (5 U/µL), 1 µL 10× Ex Taq Buffer, 0.8 µL dNTP mixture, 1 µL of each 2 µM index-primer solution (forward and reverse), 5.15 µL nuclease-free water, and 1 µL first-PCR product. Thermal cycling consisted of 94°C for 1 min followed by 10 cycles of 98°C for 10 s, 60°C for 30 s, and 72°C for 30 s.

### Standard and end-prep-omitted library preparation

Under standard conditions, second-PCR products were loaded in separate gel lanes, the target bands were excised, and the gel pieces were pooled before purification with a FastGene PCR/GEL Extraction Kit (Nippon Gene). Libraries were prepared using a Ligation Sequencing Kit V14 (ONT). The pooled amplicons underwent end preparation with a NEBNext Ultra II End Repair/dA-Tailing Module at 20°C for 5 min and 65°C for 5 min, followed by sequencing adapter ligation with T4 DNA ligase (Fig. 1a).

**Fig. 1.**
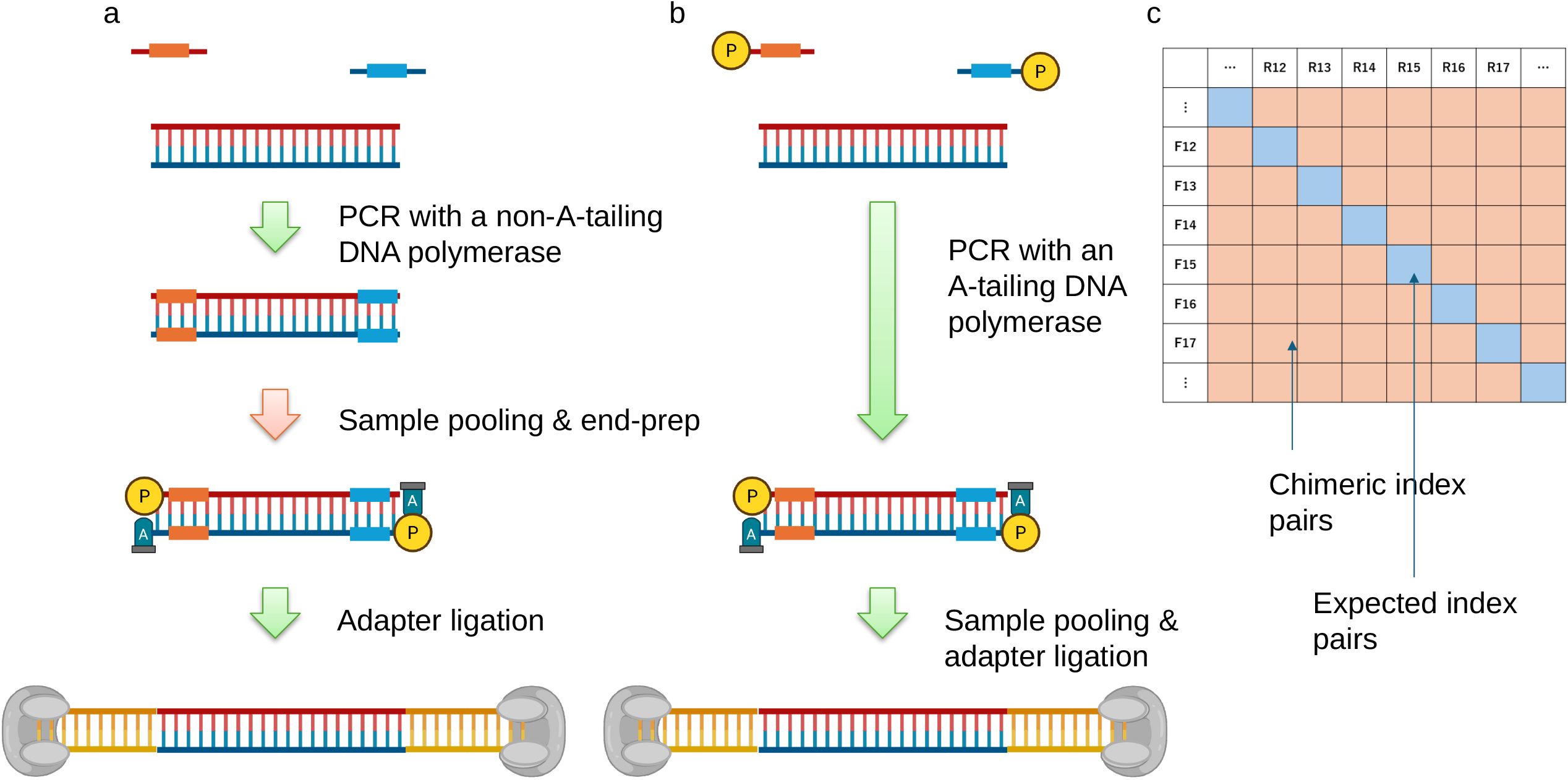
Standard and end-prep-omitted Oxford Nanopore Technologies library-preparation workflows and the unique dual-index design. (a) Standard workflow. Second-PCR products are pooled and subjected to end preparation before adapter ligation. (b) End-prep-omitted workflow. The use of 5′-phosphorylated second-PCR primers and an A-tailing DNA polymerase generates ligation-compatible amplicon ends, allowing pooled end preparation to be omitted. (c) Expected and chimeric forward–reverse index combinations. The sample-specific index pairs lie on the diagonal, whereas reads assigned to unused off-diagonal pairs are defined as index chimeras

For the end-prep-omitted protocol, the second-PCR primers were phosphorylated before use. Each 5-µL phosphorylation reaction contained 1.75 µL of 100 µM primer, 0.5 µL of 10× Kinase Buffer A, 0.5 µL of 10 mM rATP, 0.5 µL of T4 polynucleotide kinase (10 U/µL; Nippon Gene), and 1.75 µL of distilled deionized water. The reaction was incubated at 37°C for 1 h and then heated at 95°C for 5 min. PCR with the 5′-phosphorylated primers generated amplicons carrying 5′-phosphate groups and 3′-A overhangs produced by Taq DNA polymerase. The pooled end-prep step was therefore omitted, and library preparation began with sequencing adapter ligation (Fig. 1b).

Sequencing was performed using ONT Flongle R10.4.1 flow cells. Raw signal data were base-called with Dorado v1.3.1 using the super-accurate (SUP) model.

### Quantification of index chimeras

With unique dual indexing, only the forward–reverse index combinations assigned to actual samples are valid. We defined a read carrying an unused forward–reverse combination as an index chimera and used such reads as an empirical indicator of sample-index misassignment (Fig. 1c). For an unused pair consisting of forward index *x* and reverse index *y, Sx,y* was the number of reads assigned to that pair, *Fx* was the total number of reads carrying forward index *x*, and *Ry* was the total number of reads carrying reverse index *y*. The pair-specific index-chimera rate was calculated as follows:

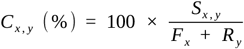

The run-level index-chimera rate was the mean of the pair-specific rates over all unused index combinations in that sequencing run.

### Dual-index demultiplexing and consensus generation

Because the barcodes varied in length and user-defined matching thresholds were required, we developed a BLAST-based dual-index demultiplexing tool, `nanopore∼split-barcode`, within PortablePipeline (https://github.com/c2997108/OpenPortablePipeline). BLAST searches were conducted using BLAST+ (Camacho et al. 2009). Demultiplexing proceeded in two stages. First, hits covering at least 60% of the full indexed-primer sequence at ≥80% identity were retained. Second, the index segment itself was required to have ≥90% coverage and ≥90% identity before a barcode was assigned. Because base calling within approximately 10 bp of ONT read termini was unstable, the primer constructs included at least 10 bp of sequence outside each index.

After demultiplexing, reads from each sample were processed in descending order of quality using VSEARCH (Rognes et al. 2016) and quality-dependent sequence-identity thresholds (Fig. 2). Reads with Q scores of 19–20 were clustered at ≥96.2% identity; reads with Q scores of 18–19 were clustered at ≥95.2%; remaining unclustered reads with Q scores of 18–20 were clustered at ≥95.2%; and reads with Q scores of 16–18 were clustered at ≥92.5%. Clusters containing at least three reads were retained, and up to 30 reads were sampled from each retained cluster and aligned with MAFFT (Katoh and Standley 2013). A majority-rule consensus was then generated at each aligned position by considering A, C, G, T, and gap states.

**Fig. 2.**
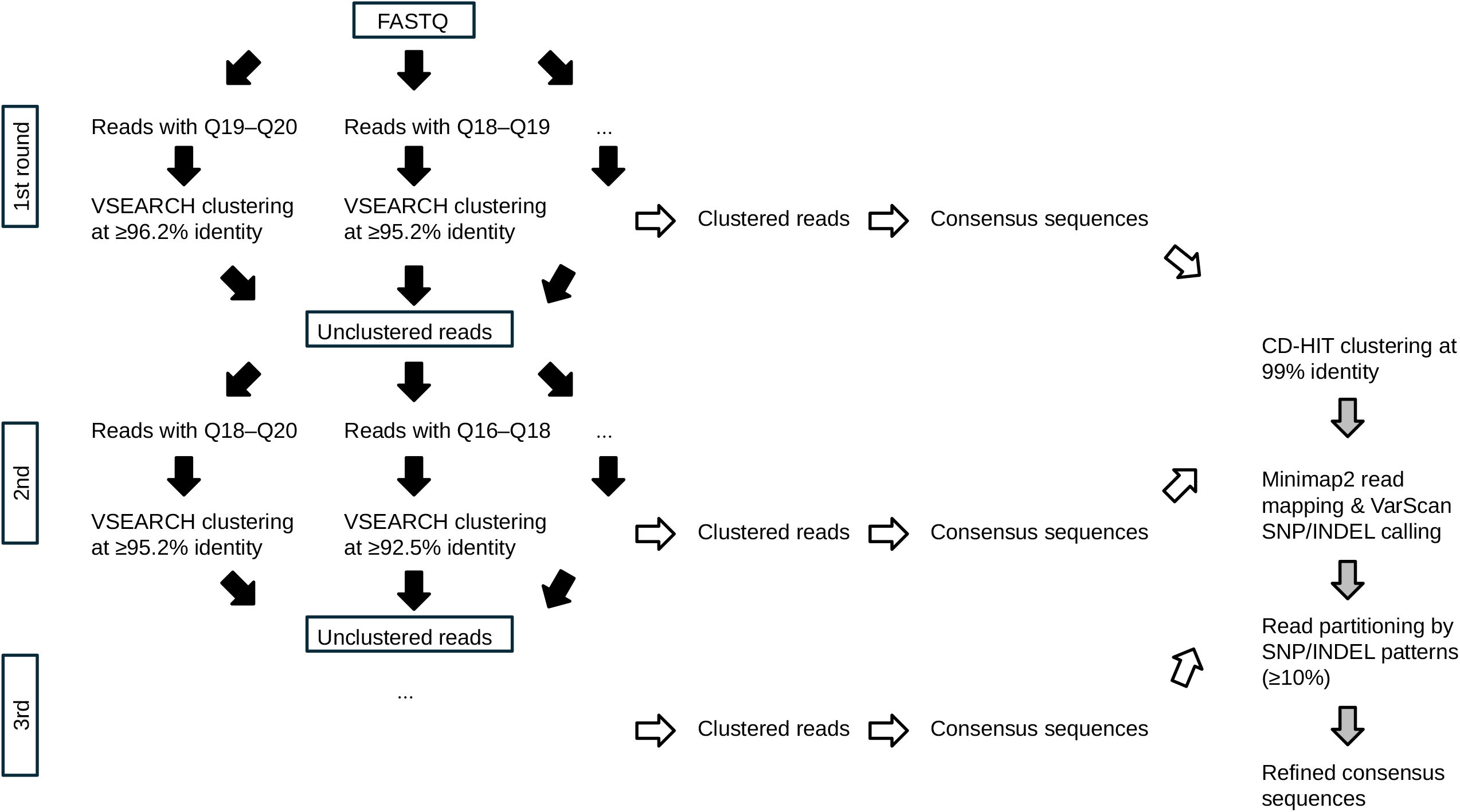
Overview of the quality-dependent consensus workflow for Oxford Nanopore reads. Demultiplexed reads are processed iteratively by Q-score bin with VSEARCH identity thresholds of 96.2% for Q19–20, 95.2% for Q18–19, 95.2% for remaining reads with Q scores of 18–20, and 92.5% for Q16–18. Clusters containing at least three reads are retained, and up to 30 reads per retained cluster are aligned to generate a majority-rule consensus. Consensus sequences are reclustered with CD-HIT at 99% identity, original reads are mapped with minimap2, SNPs and INDELs are called with VarScan, and reads are repartitioned according to SNP/INDEL patterns present in ≥10% of reads before refined consensus sequences are generated

Consensus sequences were reclustered at 99% identity with CD-HIT (Li and Godzik 2006). The original reads were mapped back to cluster representatives with minimap2 (Li 2018), and SNPs and INDELs were called with VarScan 2 (Koboldt et al. 2012). Within each cluster, reads were repartitioned according to SNP/INDEL patterns present in at least 10% of reads, and refined consensus sequences were generated. Representative sequences from all samples were dereplicated at 100% identity with CD-HIT. Finally, the original reads were reassigned to representative sequences using BLAST with ≥90% identity and a bit score of ≥150. The workflow is implemented as `nanopore∼get-consensus` in PortablePipeline.

### Taxonomic assignment and relative read abundance

Representative sequences were taxonomically assigned using metagenome∼silva-SSU-LSU_PR2_NCBI-mito-plastid_MitoFish_single-end in PortablePipeline, which performs similarity searches against public reference sequences. Operational identity thresholds were ≥99.0% for species-level assignments, ≥97.0% for genus-level assignments, and ≥95.0% for family-level assignments. Zebrafish *Danio rerio* and Japanese medaka *Oryzias latipes* were also detected in negative controls and were treated as laboratory contaminants; both taxa were removed from all ecological analyses. Table S2 contains sample-wise RRA (%) data for 180 retained taxa across 274 samples.

### Quantitative metabarcoding for comparison of archive protocols

To compare fish eDNA recovery during the Chelex and PowerSoil periods, quantitative metabarcoding was performed for samples collected in April and May using an internal standard (Ushio et al. 2018). Fifty copies of an artificial sequence were added to each first-PCR reaction. The ratio of fish reads to internal-standard reads was used to estimate fish eDNA copies per microliter of DNA extract. The artificial sequence was:

~~~
5′-
GTCGGTAAAACTCGTGCCAGCTCACCAACTGGGATGACATGGAGAAGATCTGGCACCACACCTTCTACAATGAGCTGCATGT GGCTCCCAAGGAGCACCGTATGCTGCTGACTGAGGTCCCCCTGAATCCAAGGCCAACCACAAGAAGATGACAAACTGGGATT AGATACCCCACTATG-3′
~~~

### Statistical analysis of seasonality

Seasonality was tested using the 120 samples processed with the 0.22-µm Sterivex/PowerSoil protocol from 5 April 2019 to 22 March 2022. Samples with zero total fish reads were excluded before calculating RRA. After removal of *D. rerio* and *O. latipes*, RRA values were expressed as proportions from 0 to 1 and square-root transformed. To avoid unstable tests for extremely sparse taxa, analyses were restricted to taxa detected in at least 10 of the 120 samples and in at least two of the three study years (each defined as April through the following March). Forty-six taxa passed this independent filter.

For each taxon, the harmonic regression model included study year as a fixed effect and annual and semiannual sine and cosine terms derived from calendar day. The joint contribution of the seasonal terms was evaluated using a Freedman–Lane residual-permutation test based on 9,999 permutations, with residuals from the reduced model permuted within study year (Freedman and Lane 1983). P values were adjusted across taxa with the Benjamini– Hochberg procedure (Benjamini and Hochberg 1995), and q < 0.05 was considered significant.

### Regional spawning-period information

Reported spawning periods were compiled from official online sources, all accessed on 31 August 2026: chum salmon, Hokkaido Research Organization (https://www.hro.or.jp/hro/topics/rensai/lunch/28.html); pink salmon, Hokkaido Research Organization (https://www.hro.or.jp/upload/40965/kenpou90karafuto.pdf); Pacific herring, Hokkaido Research Organization (https://www.hro.or.jp/fisheries/research/wakkanai/surveys-knowledge-of-fish/inpvt400000007sg/inpvt400000008cx.html); arabesque greenling, Hokkaido Research Organization (https://www.hro.or.jp/fisheries/research/wakkanai/surveys-knowledge-of-fish/inpvt400000007sg/inpvt40000000878.html); walleye pollock (Pacific stock), Fisheries Research and Education Agency (https://www.fra.go.jp/shigen/fisheries_resources/meeting/stock_assesment_meeting/2024/files/sa2024-sc01/fra-sa2024-sc01-02.pdf); Japanese smelt in Akkeshi, Hokkaido Research Organization (https://www.hro.or.jp/fisheries/publication/ima/o7u1kr0000004m28.html); Japanese anchovy, Hokkaido Government (https://www.pref.hokkaido.lg.jp/sr/gid/fis006.html); and Pacific cod, Hokkaido Research Organization (https://www.hro.or.jp/fisheries/research/hakodate/section/zoushoku/tpc0530000000hq8/tpc0530000000jph.html). These periods represent the typical spawning seasons in Hokkaido and do not demonstrate that spawning actually occurred at the Monbetsu sampling site.

## Results

### Reduction and residual patterns of index chimeras

Under the standard pooled end-prep condition, the mean index-chimera rate was 0.0674%. Omitting pooled end-prep reduced the rate to 0.000420%, representing a reduction of approximately 160-fold (Fig. 3a). The mean rates obtained with 10- and 24-bp indexes under the end-prep-omitted condition were 0.000522% and 0.000548%, respectively, and no clear index-length effect was observed (Fig. 3b). Many chimeric reads contained index sequences that matched valid indexes exactly, indicating that the residual off-diagonal assignments could not be explained solely by base-calling errors within the indexes.

**Fig. 3.**
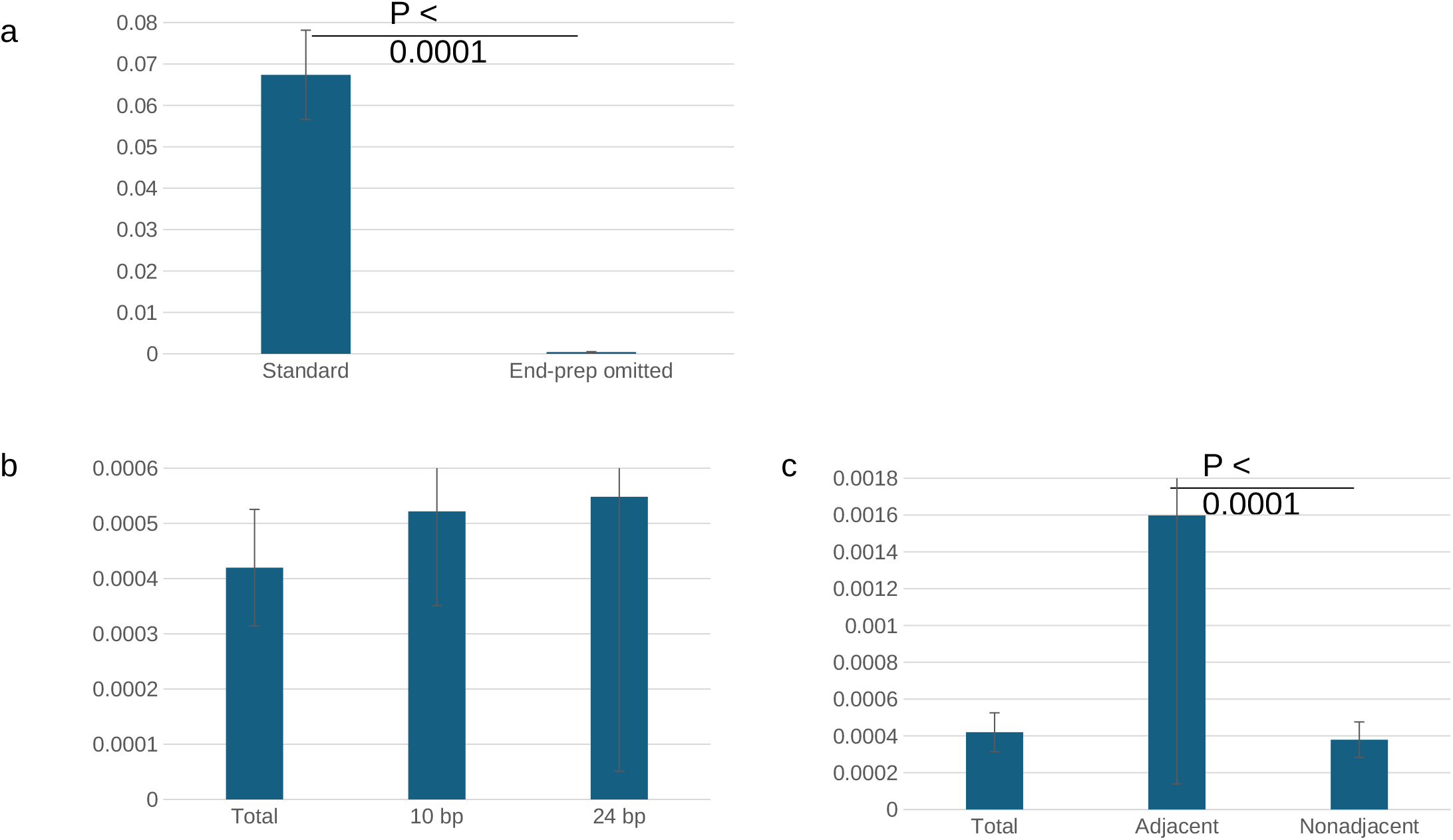
Index-chimera rates under different library-preparation and index conditions. (a) Rates under the standard protocol and the protocol in which pooled end preparation was omitted. (b) Overall rate and rates for 10- and 24-bp indexes under the end-prep-omitted condition. (c) Overall rate and rates for adjacent and nonadjacent index combinations under the end-prep-omitted condition. Error bars indicate 95% confidence intervals

The mean index-chimera rate for adjacent index combinations was 0.00160%, whereas that for nonadjacent combinations was 0.000379%, a 4.2-fold difference (Fig. 3c). Thus, the small number of chimeras remaining after omission of end-prep were not distributed uniformly among unused pairs and were enriched among adjacent combinations.

### MiFish data from the Monbetsu archive

The optimized sequencing and consensus workflow was applied to archived DNA from coastal waters off Monbetsu, yielding RRA data for 274 samples (Table S2). During the Chelex period, many extracts did not yield a visible MiFish first-PCR product, and annual amplification success rates ranged from 21.6% to 63.2%. During the Sterivex/PowerSoil period beginning in 2019, annual success rates ranged from 65.4% to 75.0% (Fig. 4a).

**Fig. 4.**
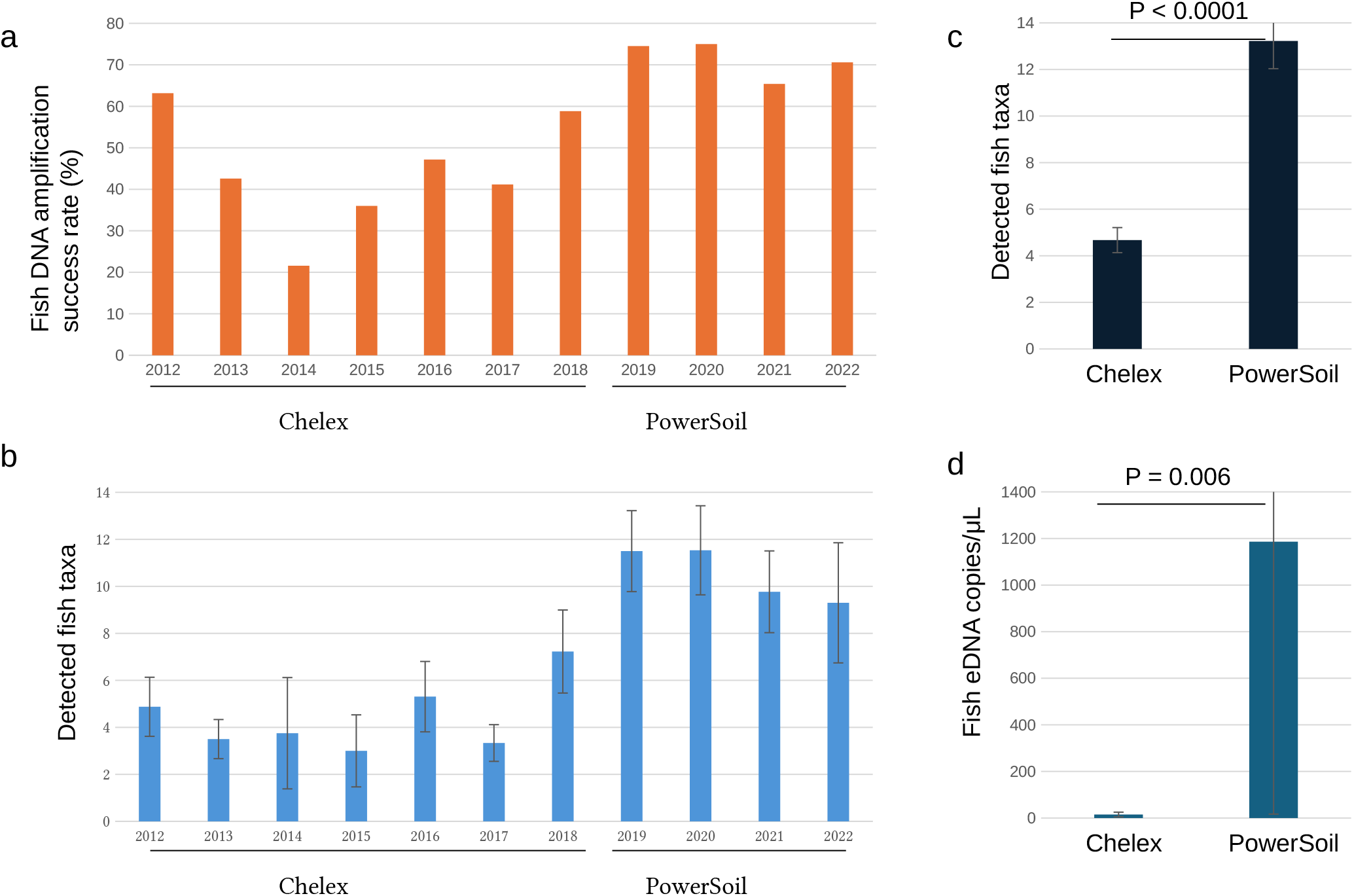
Performance of archived samples processed with different filtration and DNA-extraction protocols. (a) Annual proportion of samples in which a MiFish PCR product was obtained. (b) Annual mean number of detected fish taxa among PCR-positive samples. (c) Number of detected fish taxa among PCR-positive samples grouped according to the Chelex and PowerSoil archive periods. (d) Estimated fish eDNA copies per microliter of extract in April–May samples. The groups differ in both filtration and extraction protocols. Error bars indicate 95% confidence intervals

Among PCR-positive samples, the mean number of detected fish taxa was 4.67 during the Chelex period and 13.22 during the PowerSoil period (Fig. 4b, c). Quantitative metabarcoding of April–May samples yielded mean estimates of 14.8 fish eDNA copies/µL of extract for the Chelex period and 1,186 copies/µL for the PowerSoil period (Fig. 4d). Estimated eDNA recovery was higher with the PowerSoil method than with the Chelex method.

Monthly mean RRA values for the 20 most abundant taxa were summarized for May 2012–April 2015 (Chelex, n = 58), May 2015–March 2019 (Chelex, n = 96), and April 2019–March 2022 (PowerSoil, n = 120) (Fig. 5). Dominant taxa included Pacific herring *Clupea pallasii*, walleye pollock *Gadus chalcogrammus*, pink salmon *Oncorhynchus gorbuscha*, Japanese sardine *Sardinops sagax*, saffron cod *Eleginus gracilis*, and Japanese anchovy *Engraulis japonicus*. Because fish detection sensitivity varied among DNA extraction methods, the seasonal analysis was restricted to the 120 samples processed consistently with the PowerSoil kit.

**Fig. 5.**
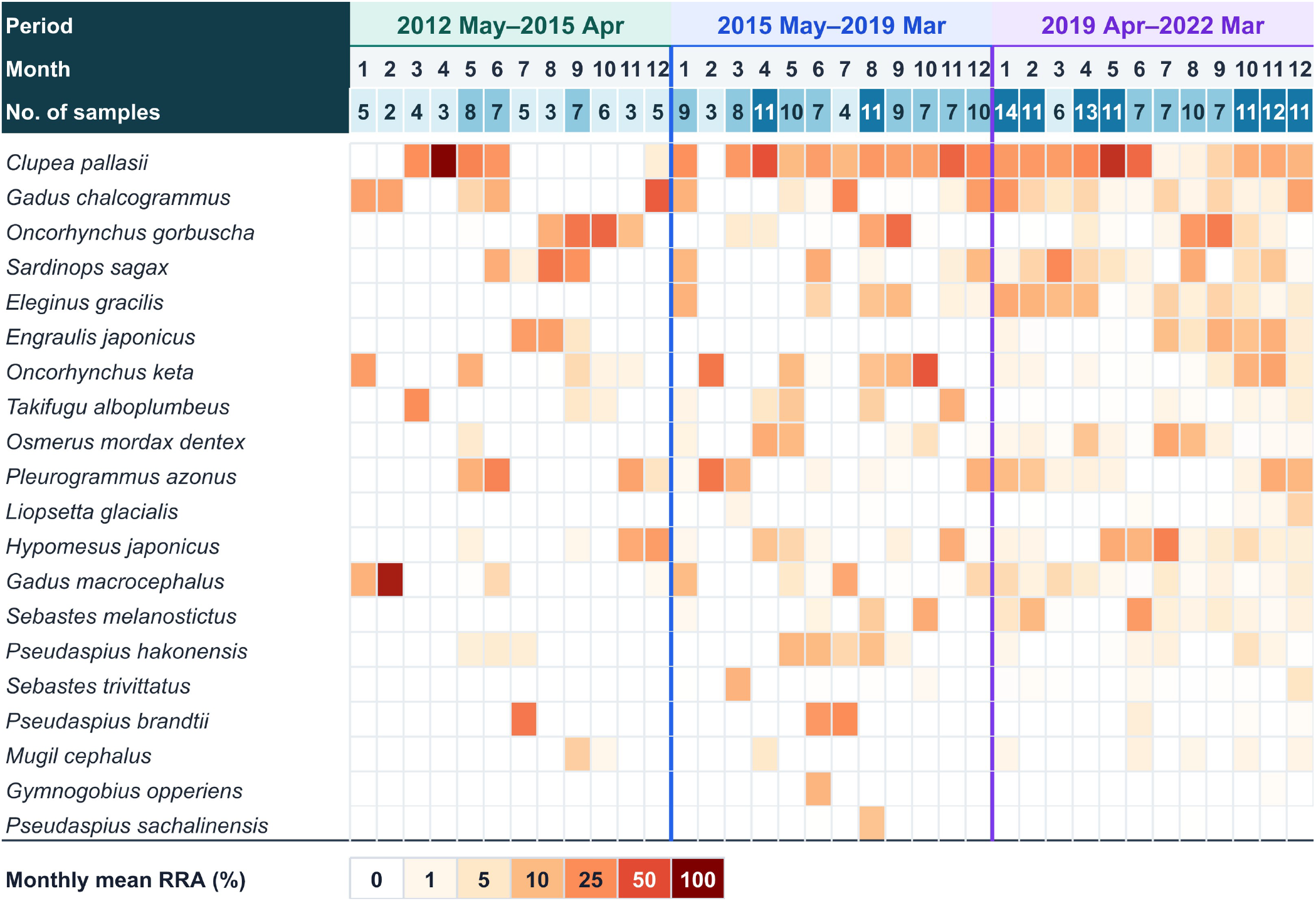
Monthly mean relative read abundance (RRA) of the 20 dominant fish taxa. Zebrafish *Danio rerio* and Japanese medaka *Oryzias latipes* were excluded. Data are shown for May 2012–April 2015 (Chelex, n = 58), May 2015–March 2019 (Chelex, n = 96), and April 2019–March 2022 (PowerSoil, n = 120). Cell shading represents monthly mean RRA (%), and the numbers above the heat map are monthly sample sizes

### Seasonal RRA patterns

Of the 46 taxa included in the seasonality tests, eight taxa had q < 0.05 and a maximum RRA >5%; regional spawning information was also available for each: chum salmon *Oncorhynchus keta*, pink salmon *Oncorhynchus gorbuscha*, Pacific herring *Clupea pallasii*, arabesque greenling *Pleurogrammus azonus*, walleye pollock *Gadus chalcogrammus*, Japanese smelt *Hypomesus japonicus*, Japanese anchovy *Engraulis japonicus*, and Pacific cod *Gadus macrocephalus* (Fig. 6). Their three-year mean RRA peaks occurred in November, August, May, November, January, July, October, and March, respectively.

**Fig. 6.**
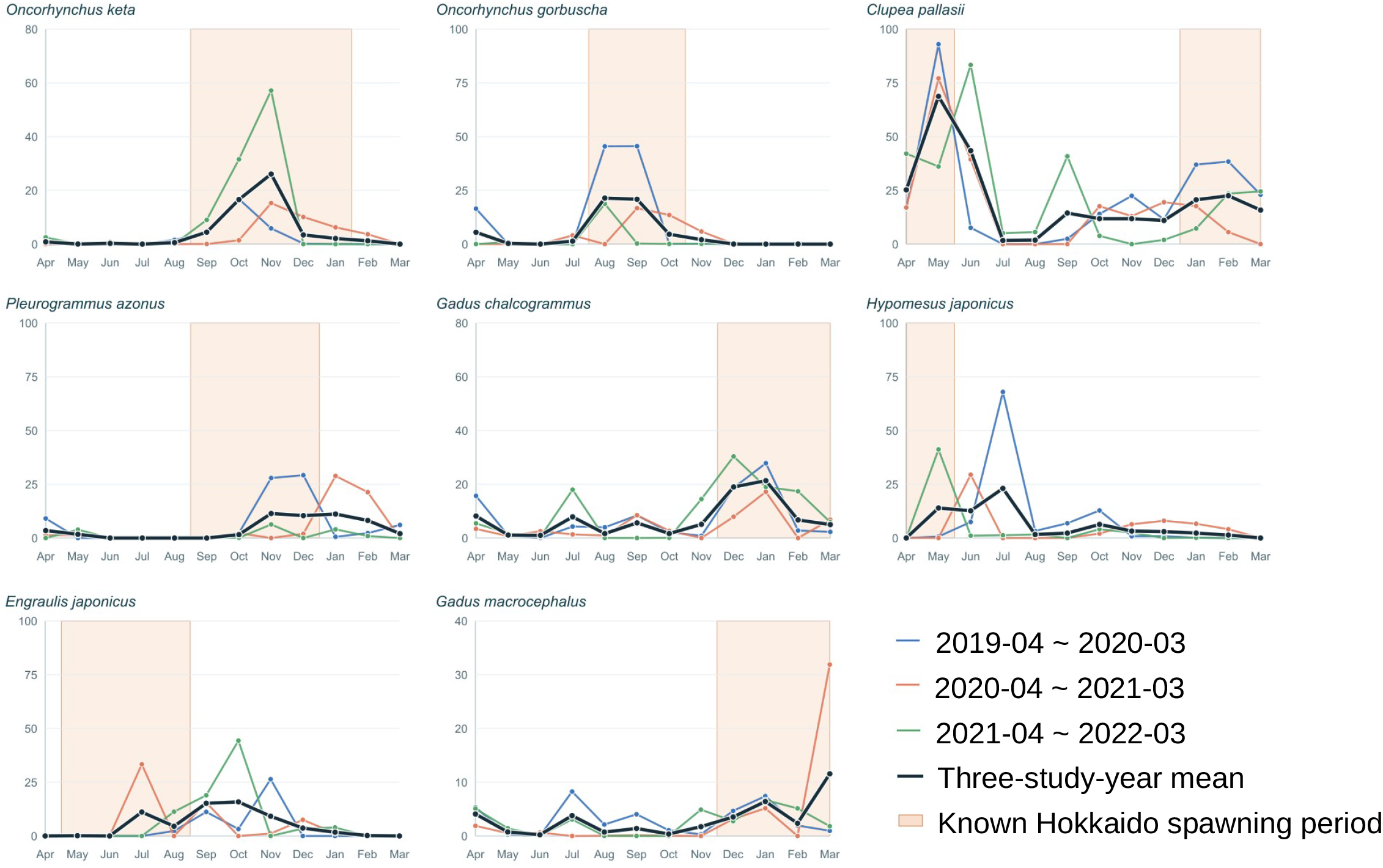
Monthly RRA of eight taxa with significant seasonal variation and reported regional spawning periods. Colored lines show monthly mean RRA for each study year from April 2019 to March 2022, and the dark line shows the three-year mean. Shaded intervals indicate reported spawning periods in Hokkaido or adjacent waters: chum salmon *Oncorhynchus keta*, September–January; pink salmon *O. gorbuscha*, August–October; Pacific herring *Clupea pallasii*, late January–early May; arabesque greenling *Pleurogrammus azonus*, mid-September–mid-December; walleye pollock *Gadus chalcogrammus* (Pacific stock), December–March; Japanese smelt *Hypomesus japonicus* in Akkeshi, late April–May; Japanese anchovy *Engraulis japonicus*, May–August; and Pacific cod *Gadus macrocephalus*, December–early March. The three-year mean RRA peaks overlapped the cited spawning periods for six taxa; Japanese smelt and Japanese anchovy peaked after the cited periods

The mean peak months for chum salmon, pink salmon, Pacific herring, arabesque greenling, walleye pollock, and Pacific cod fell within the spawning periods reported for Hokkaido (Fig. 6). The peaks for Japanese smelt and Japanese anchovy occurred after their reported regional spawning periods.

## Discussion

### Technical implications for Nanopore eDNA metabarcoding

Omission of pooled end-prep reduced the mean index-chimera rate by approximately 160-fold. Because ONT sequencing does not require clonal amplification on the flow cell, this result demonstrates that substantial index misassignment can arise outside flow-cell amplification, during or before pooled library preparation. This finding is consistent with earlier observations that post-pooling end repair or blunt ending can increase tag jumping and that residual primers and polymerase activity can promote index swapping in pooled libraries (Schnell et al. 2015; Costello et al. 2018). However, the generation of such chimeras during standard ONT library preparation has not been clearly reported previously. Applying our modified library-preparation protocol to ONT sequencing enabled us to reduce cross-sample contamination in multiplexed data to a level lower than that observed with Illumina sequencing.

The residual per-end misassignment rate of 0.000420% corresponds to 4.20 × 10^−6^ as a proportion. If misassignment events at the forward and reverse ends were independent, the probability of generating a valid but incorrect unique dual-index pair would be approximately (4.20 × 10^−6^)^2^ = 1.76 × 10^−11^, or about 1 in 5.7 × 10^10^ reads. Given that even the PromethION, ONT’s highest-throughput platform, generates on the order of 10^8^ reads, the probability of cross-sample read misassignment can be considered negligible in practice when unique dual indexing is combined with the omission of pooled end preparation.

The higher rate among adjacent index combinations further suggests that at least part of the residual signal arose before pooling or was associated with spatially structured handling. For example, low-level transfer during primer dispensing or tube/plate handling could generate such a pattern, but the present experiment did not directly identify aerosols or any other specific route. Moreover, the detection of *D. rerio* and *O. latipes* in negative controls shows that reducing index misassignment does not eliminate independent laboratory contamination. Extraction blanks, no-template PCR controls, physically separated work areas, and explicit contaminant-removal rules remain necessary.

### Consensus generation and taxonomic resolution

Consensus-based processing can substantially improve ONT metabarcoding accuracy (Baloğlu et al. 2021; Chang et al. 2024; Dubois et al. 2024). The present pipeline combines quality-dependent iterative clustering with a second partitioning step based on recurrent SNP/INDEL patterns. This design was intended to recover high-quality amplicon representatives while reducing the risk that closely related sequence variants would be collapsed into a single initial cluster. The distinct seasonal peaks recovered for *Oncorhynchus keta* and *Oncorhynchus gorbuscha* and for *Gadus chalcogrammus* and *Gadus macrocephalus* indicate that the resulting consensus sequences retained ecologically coherent species-level signals even when the MiFish region differed by only a few bases among close relatives. Although consensus generation is highly effective for improving sequence accuracy, few methods are available for reclustering reads in UMI-free Nanopore metabarcoding on the basis of SNP/INDEL variation after initial consensus generation. In particular, such methods have been reported to fail to detect taxa represented by fewer than five reads per cluster (Tedersoo et al. 2026). The analytical workflow developed in this study detects clusters supported by as few as three reads, enabling highly sensitive analysis of error-prone ONT data.

### Differences among DNA extraction methods

The optimized ONT workflow recovered interpretable long-term fish eDNA signals from DNA originally archived for plankton metagenomics. At the same time, the pronounced difference between archive periods demonstrates the importance of methodological consistency. The protocol change involved filter pore size, the number and configuration of filters, and extraction chemistry; it was also confounded with sampling year. Consequently, the higher amplification success, taxon richness, and estimated eDNA concentration during the Sterivex/PowerSoil period cannot be attributed solely to PowerSoil extraction. Long-term comparisons should therefore be based on a stable protocol or accompanied by calibration experiments in which filtration and extraction factors are varied independently.

### Seasonality and reported spawning periods

The Monbetsu time series showed strong seasonal turnover, and six of the eight focal taxa had mean RRA peaks within reported spawning periods in Hokkaido. Similar increases in eDNA during the reproductive season have been observed in freshwater and marine fishes (Bylemans et al. 2017; Takeuchi et al. 2019; Tsuji and Shibata 2021; Collins et al. 2022). Quantitative metabarcoding in a Japanese reservoir also detected eDNA anomalies that were broadly consistent with known spawning activity for many species (Wu et al. 2024). Such increases may arise from the release of gametes and associated material, from spawning migrations or aggregations that increase local biomass, or from both processes.

However, RRA is a compositional measure rather than an absolute concentration. A peak can therefore result from changes in the focal taxon, decreases in other taxa, migration, feeding aggregation, recruitment of juveniles, hydrodynamic transport, or variation in DNA persistence. The six peak–period overlaps are consequently consistent with reproduction but do not by themselves demonstrate spawning at the sampling site. For Japanese smelt and Japanese anchovy, the mean RRA peaks occurred after the cited spawning periods and may reflect one or more of these other ecological or compositional processes.

Combining absolute eDNA concentration with additional reproductive indicators could strengthen such inferences. During spawning, the release of large quantities of sperm can increase the relative contribution of nuclear DNA to the eDNA pool, thereby increasing the nuclear-to-mitochondrial eDNA ratio. (Bylemans et al. 2017; Wu et al. 2022). Integrating such markers with RRA measurements, water-temperature and current data, and fishery observations within a standardized sampling framework may enable more robust identification of spawning and recruitment events. High-frequency coastal eDNA monitoring could help elucidate seasonal patterns of nearshore habitat use.

### Conclusion

In this study, we successfully reduced index misassignment in Nanopore eDNA metabarcoding to a negligible level by combining unique dual indexing with omission of the end-prep step during library preparation. We further developed a demultiplexing and consensus-sequence generation pipeline for Nanopore data and applied it to a 10-year archive of coastal seawater samples collected off Monbetsu, revealing seasonal peaks corresponding to spawning periods. In the future, protocol standardization could enable the low-cost analysis of large numbers of samples from multisite time series, thereby supporting predictions of ecosystem responses to climate change.

## Declarations

### Author contributions

Masatoshi Endo and Noriko Kurita performed the PCR experiments. Satoshi Nagai conducted sample collection and DNA extraction. Tsuyoshi Watanabe performed DNA extraction. Shuichi Asakawa contributed to the scientific discussion and interpretation of the results. Kazutoshi Yoshitake conceived and coordinated the overall study and wrote the manuscript.

### Data and code availability

Unique dual-index primer sequences and sample-wise RRA data are provided in Tables S1 and S2. The raw Oxford Nanopore sequencing data have been deposited in the DNA Data Bank of Japan (DDBJ) under BioProject accession no. PRJDB43058. The demultiplexing and consensus workflows are available in PortablePipeline at https://github.com/c2997108/OpenPortablePipeline.

### Ethics approval

Not applicable. The study analyzed archived seawater DNA and did not involve the handling of live vertebrate animals.

## Electronic supplementary material captions

**Table S1** Unique dual-index primers used in the second PCR. Primer name, direction, ID, index length, index sequence, and full primer sequence are provided for 60 forward and 60 reverse primers. All sequences are shown in the 5′–3′ direction

**Table S2** Sample-wise relative read abundance of fish taxa after exclusion of two laboratory contaminants. The matrix contains 180 taxa and 274 samples. Values are RRA (%). The remaining values were not renormalized after removal of *Danio rerio* and *Oryzias latipes*

